# Variance spectrum scaling analysis for high-dimensional coordination in human movement

**DOI:** 10.1101/2025.01.07.631755

**Authors:** Dobromir Dotov, Jingxian Gu, Philip Hotor, Joanna Spyra

## Abstract

Coordinated movement has been essential to our evolution as a species but its study has been limited by ideas largely developed for one-dimensional data. The field is poised for a change by high-density recording tools and the popularity of data sharing. New ideas are needed to revive classical theoretical questions such as the organization of the highly redundant biomechanical degrees of freedom and the optimal distribution of variability for efficiency and adaptiveness. Methods have been focused on increasing dimensions: making inferences from one or few measured dimensions about the properties of a higher dimensional system. The opposite problem is to record 100+ kinematic degrees of freedom and make inferences about properties of the embedded manifold. We present an approach to quantify the smoothness and degree to which the manifold is distributed among embedding dimensions. The principal components of embedding dimensions are rank-ordered by variance. The power-law scaling exponent of this variance spectrum is a function of the smoothness and dimensionality of the embedded manifold. It defines a threshold value beyond which the manifold becomes non-differentiable. We verified this approach by showing that the Kuramoto model close to global synchronization in the upper critical end of the coupling parameter obeys the threshold. Next, we tested if the scaling exponent was sensitive to participants’ gait impairment in a full-body motion capture dataset containing short gait trials. Variance scaling was highest in the healthy individuals, followed by osteoarthritis patients after hip-replacement, and lastly, the same patients pre-surgery. Thinking about manifold dimensionality, smoothness, and scaling could inform classic problems in movement science and exploration of the biomechanics of full-body action.

## 1. Introduction

Stable and efficient movement over various terrains is a skill that was necessary for our survival as a species throughout the evolutionary time scale, and is essential for independent daily living on the time scale of individual lifespan. To achieve upright bipedal locomotion, the human body needs to organize a system consisting of many bio-mechanical degrees of freedom. Traditionally referred to as Bernstein’s degrees of freedom question (foundational in human movement science, motor control, and related branches of neuroscience ^1^, the challenge is to understand how the nervous system resolves the redundancy of biomechanical degrees of freedom while ensuring context-sensitivity to environmental and task constraints ^2^. Presumably, it does this by exploiting a hierarchical approach to organization ^3^. A more recent approach has been to reframe the redundancy as a bliss of motor abundance because spare resources can be used to achieve adaptive properties such as stability, efficiency, and resistance to perturbations ^4,5^.

This theoretical problem can be illustrated in terms of its implications for the acquisition of motor skills ^6^. For example, a beneficial strategy for beginning to learn how to throw a basketball appears to be to lock all degrees of freedom but one and use it to complete the desired trajectory. This is the *freezing* stage of learning. With continued learning, one begins to unlock degrees of freedom so that they make complementary contributions to the throw. This is the *release stage*. To achieve skilled performance, players are often instructed that the shot should fluidly progress from the feet all the way to the wrist and fingers. Indeed, evidence confirms that center of mass (related to body’s posture) and ball release variables (how and when the ball leaves the hand) are coordinated but not locked to each other in skilled throws ^7^. Despite its intuitive nature, however, the freeze-release principle of motor learning is not supported conclusively by the overall evidence in the literature ^8^. This is partly due to the historical emphasis on relying on a small number of focal kinematic and kinetic variables instead of comprehensively analyzing full-body movement.

It is a remarkable fact that little effort has been made to understand full-body coordination quantitatively despite the foundational and still unsolved status of the degrees of freedom problem, the importance of hierarchical control, and the ambiguity in associated problems such as coordination during development and skill acquisition. One historical reason for this blind spot of the movement sciences has been the complexity and cost of high-density motion tracking techniques. More recently, various forms of motion tracking technology, including markerless motion tracking, have become practical. Furthermore, the ubiquity of data sharing practices has made it cost-effective to apply different computational techniques to publicly available rich datasets, an approach that has enabled tremendous progress in other domains such as machine learning and AI. The time is ripe to breathe new life in classical theoretical questions in movement science.

The remaining obstacle in this context has been the lack of analytical tools, with few exceptions such as the human movement kinectome ^9^. The predominant multivariate tools tend to be extensions of uni- and bi-variate methods for coordination dynamics. It is unlikely that these will scale to dozens or even hundreds of variables because of computational and theoretical limitations. Any method that begins by parameterizing each pairwise relationship will have to exhaust all possible pairs. The number of these pairs, and the corresponding computational time and parameter space, accelerates with the number of dimensions. The family of methods based on recurrence quantification analysis, popular in the movement sciences, have been extended first to two, and then to more variables ^10^. This comes at a cost, though, as it requires estimating parameters for each input variable. Furthermore, simply extending a method to receive more variables does not guarantee that it will also extend its interpretation. For example, the order parameter *R* generalizes pairwise relative phase for collectives of oscillators, yet it fails to distinguish between scenarios of uniformly poor synchronization among all oscillators and scenarios of strong but highly clustered synchronization. In the following, we propose an interpretable and inherently multivariate analysis of high-density coordination.

### 1.1. From dimensionality reduction to variance spectrum scaling

Here, we propose a method for measuring and interpreting geometrical properties of manifolds embedded in high-dimensional time-series data. What we mean by manifold is a *d*-dimensional surface that curves and loops on itself as it is embedded in an *n*-dimensional observation space. A typical first step when dealing with such data is to apply a dimensionality reduction method to remove redundancies shared between variables. Indeed, principal component analysis (PCA) can be used in the study of full-body movement kinematics to help focus analysis on relevant patterns of movement ^11,12^. PCA is a dimensionality reduction technique that projects a given dataset into a new coordinate system such that each axis, called principal component (PC), explains as much variance of the original data as possible while staying orthogonal to the other axes. Intuitively, PCA can be thought of as taking a camera that is pointing at a cloud of data points and turning it in such a way that the data line up in a useful axis, like looking at a tree from the top versus the side. Dimensions containing matching correlated patterns line up relative to the camera and are seen as one, and unique patterns are made more visible by projecting them on unique axes to emphasize their variance. Importantly, PC’s are ranked by the amount of variance they explain. In this way, the first PC’s contain gross features of the data and subsequent PC’s contain ever finer detail until only noise is left. PCA has been widely used in multiple fields as a way of compressing information by retaining only the top PC’s, for testing to what extent information is subject to compression, or for emphasizing unique information.

It is less often appreciated that PCA can be used not only for reducing dimensionality but also for measuring geometric properties of the lower-dimensional manifold ^13^. The rate at which the variance (explanatory power) decreases over consecutive PC’s is related to the fractal geometry of the manifold on which the original data live. The decay rate of the so-called variance spectrum or eigenspectrum is bounded by the smoothness and dimensionality—independently—of the unique manifold embedded in the high-dimensional observation space. The decay of the variance spectrum is given by the following scaling law,

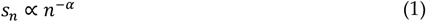

If the original data live on a smooth manifold, meaning that the manifold is differentiable everywhere, then its variance spectrum must decay faster than or equal to a threshold

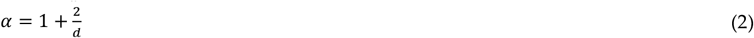

where *d* is the dimension of the manifold, see Figure 1d-f. Note that for *d* → ∞ this becomes the familiar power-law *s* ∝ *n*^−1^ arising in various forms in one-dimensional observations. Its implications for higher-dimensional problems are yet to be investigated.

**Figure 1.**
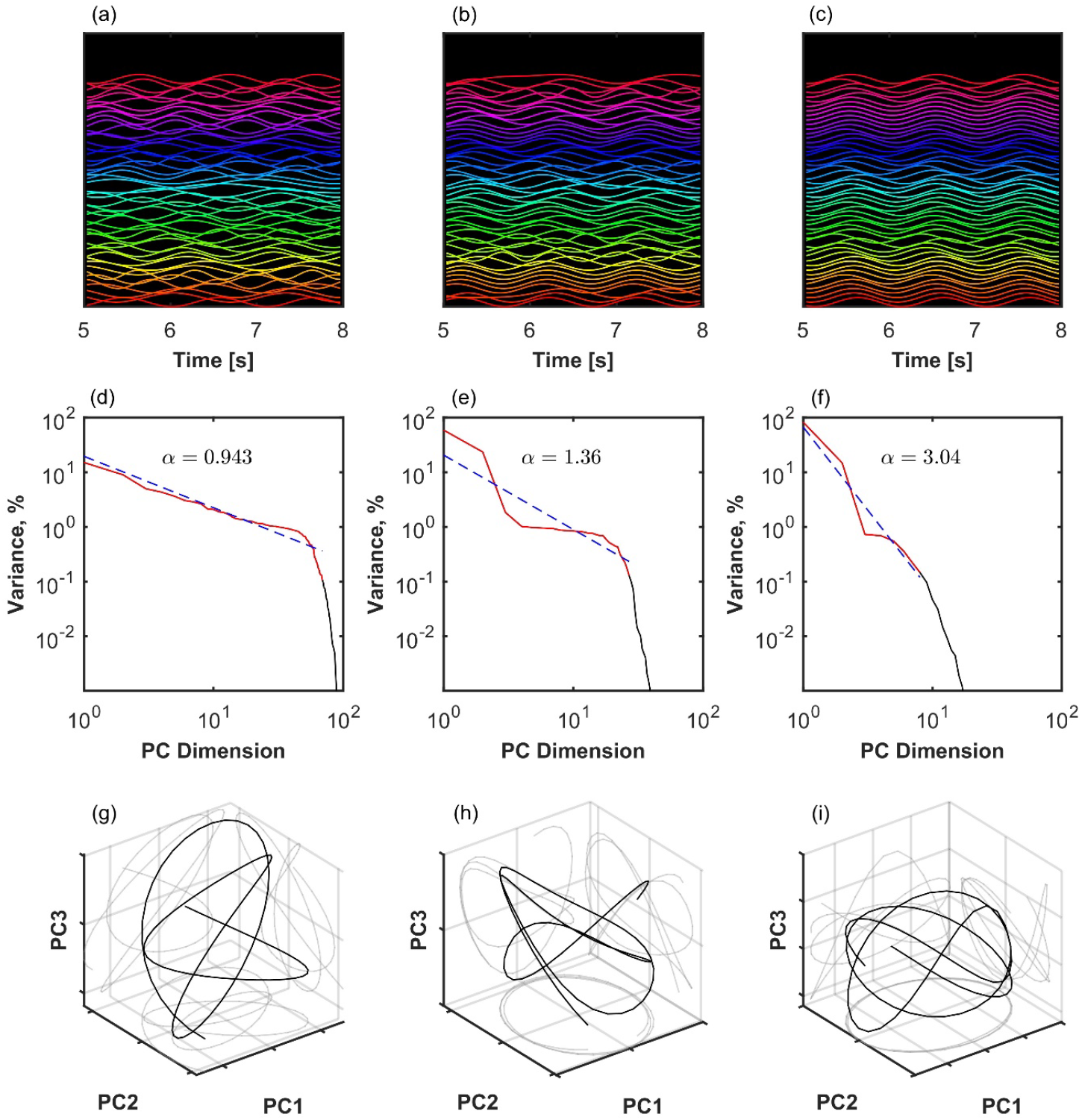
Variance spectrum analysis applied to three examples of the finite size Kuramoto system with *N*=100 oscillators and different levels of coupling. Left column (a, d, and g): Weak coupling at the beginning of the critical region, *K*=2. Middle column (b, e, h): Moderate coupling in the middle of the critical region, *K*=2.7. Right (c, f, and i): Strong coupling close to global synchronization, *K*=4. (a-c): Time series of simulated oscillators. A sparse sample of oscillators shifted on the *y*-axis and in a short time window are shown for visibility. (d-f): Eigenspectrum scaling computed by fitting an inverse power-law *s*_*n*_ ∝ *n*^−*α*^ (red line) to the variance of principal components (black). In the first two columns (a,d,g and b,e,h), the observed scaling was *α* < 3, consistent with lack of global synchronization and manifold *d* > 1, see Eq. 2. (g-i): The first three principal component projections of the oscillator collectives.

The relationship between observation space, scaling of the principal component variance spectrum, manifold dimension, and manifold differentiability was explained in the context of information encoding in neural space consisting of thousands of neurons ^13^. Here we will give it a biomechanical interpretation. To improve the coding efficiency of motor control, it would be convenient to distribute separate aspects of the movement to independent degrees of freedom, as in early robotics designs with no redundancy among the joints and motors ^14^. In variance spectrum space, this would result in a low α: many small principal components with flat eigenvalues. This would likely violate the manifold smoothness condition, as the trajectory would be jumping discretely among dimensions of the manifold. On the opposite end, control could be simplified by locking degrees of freedom to each other, the freeze stage of learning. In variance spectrum space, this would result in high α: few principal components account for all variance and eigenvalues decay very fast. Our hypothesis is that biomechanical systems are positioned above but close to the critical scaling 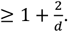

In this range, the underlying manifold is differentiable, hence movement is efficiently smooth, but still distributed among as many degrees of freedom as possible to improve stability by allowing degrees of freedom to absorb perturbations and complement each other. In the following section, we illustrate these ideas using simulated Kuramoto systems of oscillators as minimal examples of systems with phase-locking among its variables. Then we test the hypothesis by analyzing full-body kinematics during walking recorded in human participants with varying levels of motor disability.

## 2. Materials and Methods

### 2.1. Finite size Kuramoto system around the critical region of coupling strength

The Kuramoto dynamic system of coupled phase oscillators, Eq. 3, was conceptualized as a model of coupling-dependent spontaneous transition to a globally ordered state exhibited by populations of dynamic units with different natural frequencies

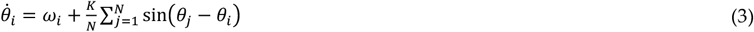

Here, θ_*i*_is the phase of oscillator *i*, each has a preferred frequency ω_*i*_ describing how fast around the unit circle it likes to go, *K* is coupling strength, and *N* is number of oscillators. Finite size and finite time simulations do not necessarily reflect the stable analytical properties of the ideal model ^15,16^. Nevertheless, they exhibit their own interesting dynamic phenomena ^17,18^, and have provided model-based insight about synchronization in a large variety of natural systems (for some examples, ^19–21^).

The model is suitable for testing the relationship between redundancy, manifold dimension, and variance scaling in a system of dynamic variables with varying synchronization, ranging from total independence to total convergence to a smooth global manifold with *d* = 1. This is because the model gives a mathematical account of how individual oscillators become enslaved by the mean field. This can be seen by using the definition of the mean field of phases, Eq. 4, to express Eq. 3 equivalently in terms of the coupling between individual oscillators and the mean field, Eq. 5. ψ is the mean field phase. Global synchronization, the coherence among oscillators, is measured by the mean field amplitude *r*, also called order parameter.

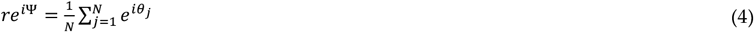

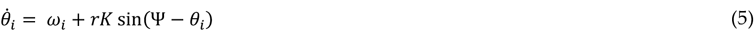

We assumed that the oscillators are also subject to Gaussian random variation 𝒩(0, *σ*).

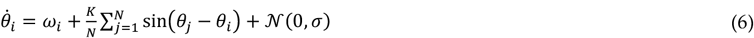

To keep the system commensurate in size with the empirical gait dataset presented below, we simulated Eq. 6 with *N* = 100, *σ* = 2 over 100 trials and varied *K* ∈ [0,8] uniformly across trials, see Figure 2a. Each trial, the distribution of intrinsic frequencies ω_*i*_was drawn from a Gaussian distribution 𝒩(0, *γ*) with *γ* = .25. In each trial had a simulated duration of 100 *s* at a sampling rate of 300 Hz. We used a third-order Runge-Kutta solver. Note that the theoretical critical *K* cannot be determined exactly in a finite size model.

**Figure 2.**
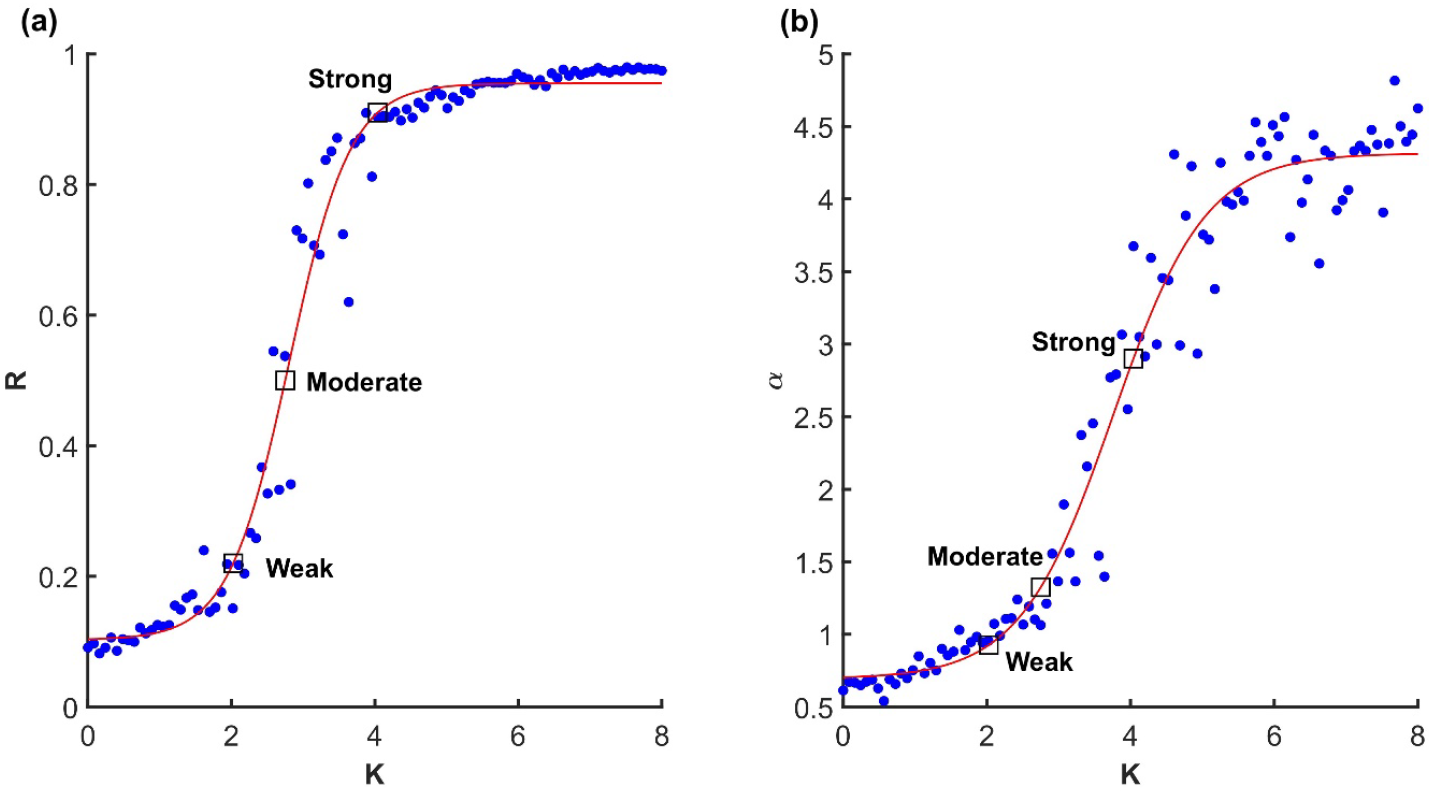
(a) Relationship between coupling strength *K* and synchronization (order parameter) *R* in the finite size Kuramoto system with *N*=100. (b) Relationship between coupling strength and variance scaling. The squares correspond to the three representative scenarios illustrated in Figure 1. Dots are trials and red lines are fitted sigmoid functions.

#### 2.1.1. Analysis

PCA was performed on the real component of the phase oscillators. We used singular value decomposition after zero-centering each variable. The eigenvalues of each principal component were converted to variances in the range from 0 to 100 percent. We fitted a power law of the form *s*_*n*_ ∝ *n*^−*α*^ (Eq. 1) to the rank-ordered variances, see Figure 1d-f. To remove the drop off region we discarded variances below a fixed threshold of .1%. We also fitted a Zipf-Mandelbrot power law which has two more parameters, but it appeared not to fit the curves as well.

### 2.2. Human gait recorded with high-density motion tracking

For the human movement data, we used a publicly share dataset ^22^ with full-body motion capture of participants performing short walking trials (six meters straight line).

#### 2.2.1. Study population

The participants included healthy adults and patients with unilateral hip osteoarthritis (OA). Patients were recorded twice, before hip-replacement surgery and six months after successful surgery. The asymptomatic group included 80 healthy participants between the ages of 25 and 82. There were 106 patients with hip OA without other diseases between the ages of 45 and 85.

#### 2.2.2. Protocol and equipment

Participant walked back and forth a six meters straight line at a self-paced speed. Each condition was recorded for a minimum of ten trails. Eight optoelectronic cameras sampled at 100Hz (Vicon MXT40, Vicon, UK) and 35 reflective cutaneous markers placed on anatomical landmark locations were used to record 3D kinematics, resulting in a total of 105 position time-series variables.

#### 2.2.3. Analysis

For pre-processing, data quality was controlled by removing bad trials with markers missing for more than 50% of the duration of the trial. The positional drift due to forward translation while walking was removed by regressing out the translation. To equalize data length across trials, participants, and groups, each trial was re-sampled to 500 time points using linear interpolation, so that there were roughly 100 samples per second.

The analysis followed the procedure described in Section 2.1.1. The only difference was that the cutoff threshold was set to a lower value, .01%.

## 3. Results

### 3.1. Variance scaling in the Kuramoto system

To confirm that the Kuramoto model was valid, and that the critical transition region of parameter space was located, we first focused on synchronization as a function of coupling strength, Figure 2a. Indeed, the relationship between the order parameter *R* and coupling strength *K* exhibited the familiar sigmoid shape although not with a sharp transition from 0 to 1 as expected from the ideal model.

Next, we consider variance scaling at selected points in parameter space. As expected, we found that when the model was in the higher end of the critical region, variance scaling crossed *α* = 3, Figure 2a-b, which is the threshold of smoothness for *d* = 1, see Eq. 2. We assumed that manifold was *d* = 1 because, by definition, the Kuramoto model in its coherent state attracts all units to one synchronized oscillator. In short, when coupling is set just above the critical value, the Kuramoto model exhibits critical balance between globally synchronized and smooth manifold and maximum possible distribution of variability among its embedding dimensions.

### 3.2. Variance scaling in human gait

The analysis of representative trials from each group is illustrated in Figure 3. Statistical analysis with a linear mixed-effects model showed that healthy participants exhibited the highest scaling, consistent with strongest synchronization among kinematic degrees of freedom, followed by patients after successful surgery (β = −0.318, *SE* = 0.047, *p* < 0.001, Cohen’s *d* = 0.680). Scaling in patients before arthroplasty surgery was lower than after surgery (β = −0.108, *SE* = 0.013, *p* < 0.001, Cohen’s *d* = 0.230). As expected, all groups were close to but above the critical value of *α* = 3 on average, see Figure 4. Interestingly, the pre-surgery group was closest to the critical value.

**Figure 3.**
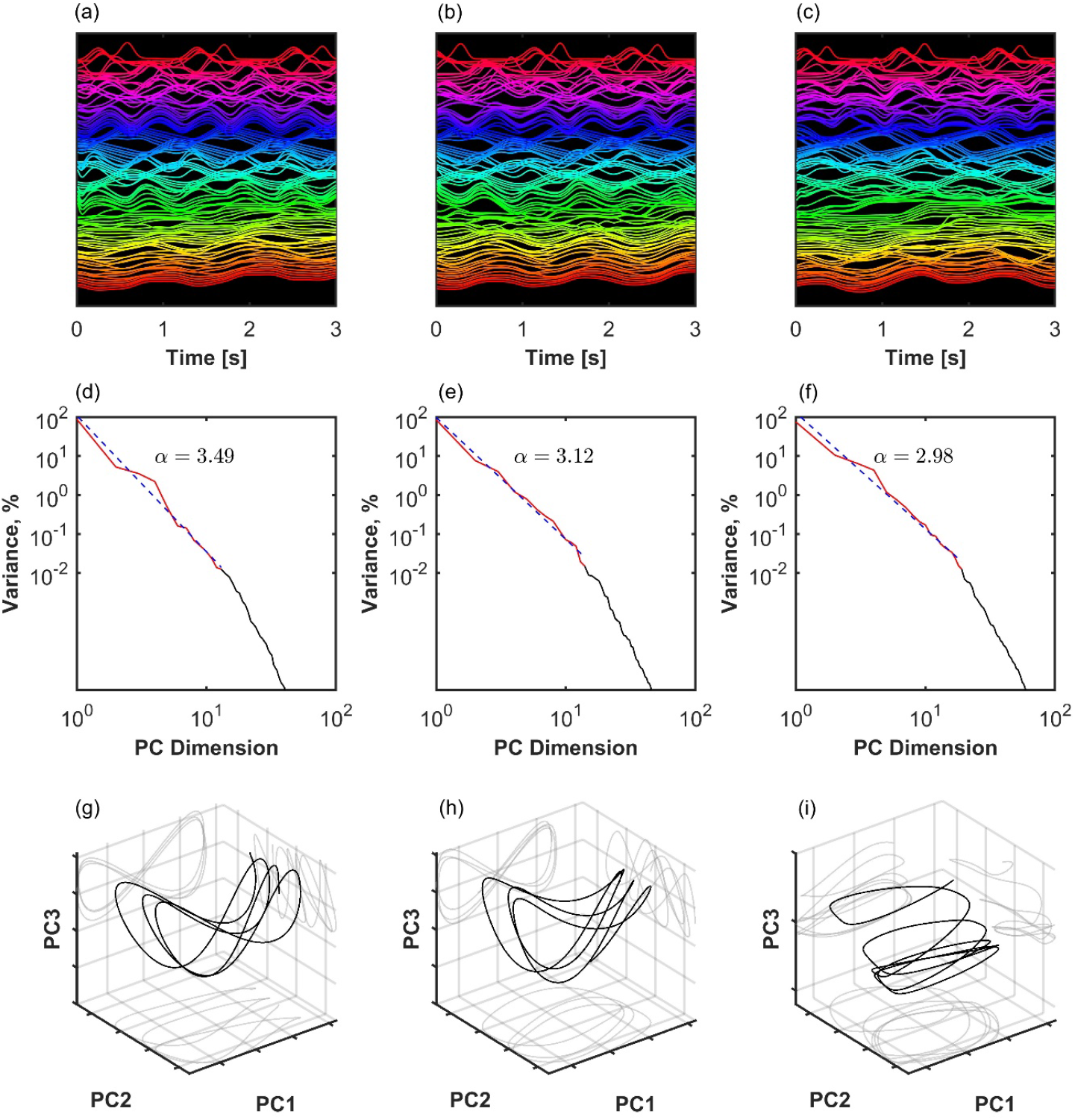
Three examples of coordination analysis in the human kinematic gait data with *N*=105 markers. (a-c): Time series of kinematic markers. For better visibility, a short time window is shown, and time series are normalized with the z-score and shifted on the y-axis. (d-f): Eigenspectrum scaling computed by fitting an inverse power-law *s*_*n*_ ∝ *n*^−*α*^ to the variance of principal components. (g-i): The first three principal component projections. Left column (a,d,g): Healthy participant with a high scaling exponent. Middle column (b,e,h): Patient after successful arthroplasty exhibits slower decay of the variance spectrum, meaning that there is more variance distributed among the less important dimensions of the manifold. Right column (c,f,i): Patient before arthroplasty exhibits even slower decay of the variance spectrum. In the latter case, scaling approaches the theoretical lower bound for a smooth manifold 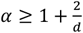 for *d*=1.

**Figure 4.**
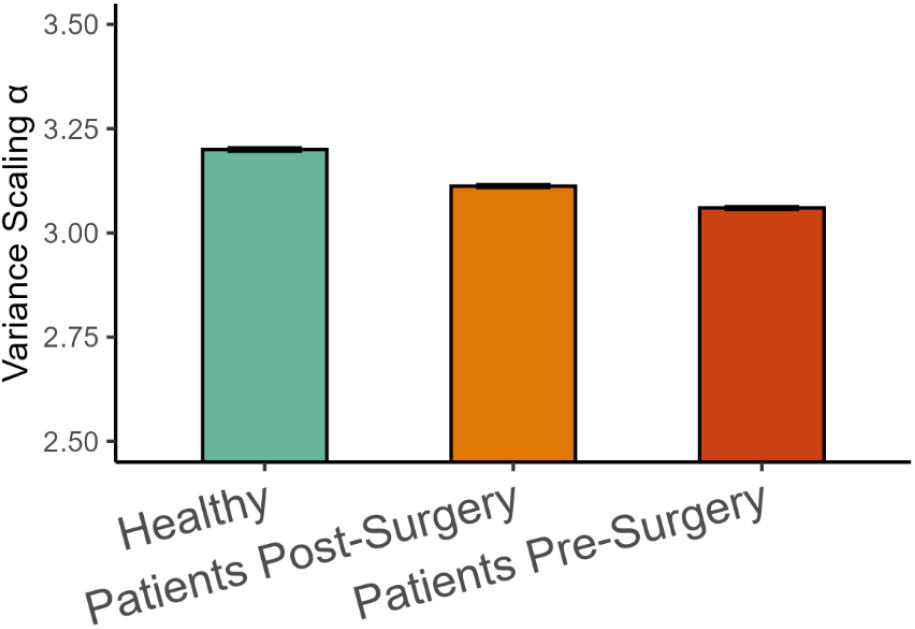
Variance scaling exponents (Mean +/− 95%CI) in the three groups of participants.

## 4. Discussion

We introduced an approach to investigating properties of full-body movement manifolds as they are embedded in much higher dimensional observation spaces comprising potentially hundreds of kinematic markers. The rate of decay of the eigenspectrum (variance spectrum) after dimensionality reduction is related to the smoothness of the manifold and the distribution of coordination variability among different degrees of freedom. We defined a reference value, Eq. 2, that enables principled interpretation of the observed results in terms of boundary and optimality conditions. The ideas were confirmed using the Kuramoto model, a system of phase oscillators with various amounts of phase-locking between them and a critical region of coupling strength. We observed that the super-critical regime was associated with optimal balance between smoothness of the manifold of the globally synchronized system and distribution of variance among the dynamic units.

We then studied whether full-body movement organization during walking was characterized by a similar critical regime. The evidence suggests that healthy walking was in the super-critical regime for a low dimensional task and that disability was associated with lowering the scaling value closer to the critical threshold, therefore releasing degrees of freedom but also potentially leading to discontinuities in the manifold derivative.

This is a minimally presumptive approach to investigating high-density coordination and it has the potential to inform classic issues in movement science. Elsewhere, the question of task and neural space dimensionality has generated a lively debate concerning the structure of the neural substrates supporting different activities ^23–31^. In the context of neural coding, an efficient approach that preserves resources is one where information is uniquely coded in separate subspaces. On the contrary, a stable approach is one in which all degrees of freedom are involved redundantly as a neural population.

A similar question concerns the structure of multi-dimensional biomechanical variability supporting different tasks ^32^. Historically, progress has been limited by expensive methods for comprehensive tracking and analysis of movement. In fact, the movement sciences often deal with the problem of up-dimensionalizing few observed variables to uncover inherent system dynamics. A popular method for this is phase space reconstruction ^33^. In contrast, we are now facing the opposite problem: high-density and highly redundant datasets require dimensionality reduction techniques to focus on the relevant system dynamics.

To help interpret the geometry of full-body dynamics embedded in many motor degrees of freedom, we pointed out that the manifold distribution and smoothness serve to define a reference scaling value by combining two constraints with opposite directions. This is based on biomechanical considerations. On one hand, the smoothness condition considers the fact that sharp transitions in the manifold of positional coordinates would imply diverging velocities. It would be an inefficient control strategy to correct full-body movement by sharp sudden impacts, not to mention that infinite velocity is hard to make sense of in a mechanical system. Mathematically speaking, non-differentiable gait cycle dynamics imply instability, falls, and, generally, unpredictability. This is a tendency to increase the absolute slope of the singularity spectrum. On the other hand, distributing variability across a wider range of degrees of freedom is a strategy for flexibility and resilience to perturbations. This is a tendency to decrease the absolute slope of the singularity spectrum. The combination of both tendencies leads to optimal scaling in the vicinity of the smoothness threshold 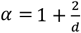.

Here we assumed that the task of walking is fundamentally one dimensional but distributed among degrees of freedom. This can be motivated by the theoretical debate whether bipedal gait can be reduced to a simple one-dimensional template model ^34^ such as inverted pendulum ^35,36^, swinging limb as a pendulum ^37^, etc.. Plugging *d*=1 in Eq. 2 led to *α* = 3 which is in remarkable agreement with the empirical observation. It is possible, however, that the assumption oversimplifies gait dynamics. Further research is necessary to determine if the threshold exponent is not lower because the manifold is larger and that other factors do not contribute to the optimal smoothness and distribution of variability across embedding degrees of freedom.

Another theoretical advantage of addressing full-body coordination is that it avoids relying on weakly justified choices about which coordination pair is the relevant one. In some instances, relevance is implied by the task, as in swinging limbs to maintain a given rhythm for ambulation ^38,39^. Research in human coordination dynamics has tended to focus on experimentally constrained tasks with isolated coordination between two effectors, be it fingers or limbs. It has paid less attention to how the task is supported by full-body dynamics in the background.

The present approach has further potential to inform classical ideas in movement science. The objective of exploring all dimension of variability, whether they are obviously involved in the task or not, is mathematically related to the highly influential (un)controlled manifold framework ^40,41^. Furthermore, we are yet to explore whether the optimal properties of the high-dimensional manifold relate to the ideas of optimal variability and so-called biological variability of one dimensional movement variables ^42,43^.

## Institutional Review Board Statement

The openly accessible gait dataset was collected in accordance with the Declaration of Helsinki and the Good Clinical Practice (ICH Harmonised Tripartite Guideline, 1996) and approved by the local ethic committee (CPP Est I, Dijon, France).

## Data Availability Statement

No new data were created.

